# Proteome analyses reveal Endoplasmic Reticulum stress-induced changes in protein abundance associated with Ube2j2 deficiency in human cell culture

**DOI:** 10.64898/2026.03.31.715661

**Authors:** C. L. Dahlberg, M. Zinkgraf, S.H. Laugesen, C.L. Søltoft, Q. Ginebra, E.P. Bennett, R. Hartmann-Petersen, L. Ellgaard

## Abstract

The unfolded protein response (UPR) helps reinstate cellular proteostasis upon an accumulation of misfolded proteins in the endoplasmic reticulum (ER), in part through ER-associated degradation (ERAD). Ube2j2 is an ER-localized E2 ubiquitin-conjugating enzyme that participates in ERAD. We used mass spectrometry analysis of cultured U2OS cells to investigate how the loss of Ube2j2 affects the cellular proteome in response to tunicamycin-induced ER stress. We constructed a network of twelve statistically distinct modules of protein abundance profiles across conditions. We describe the Gene Ontology annotations for each module along with the “hub gene” proteins whose abundance levels most closely adhere to each module’s protein abundance profile. Our analysis identifies known Ube2j2-associated pathways (e.g., the UPR and ERAD) and cellular functions that were previously unassociated with Ube2j2 (e.g., RNA metabolism, ER-Golgi transport, and cell-cycle progression). These data are available via ProteomeXchange with identifier PXD076153 and provide avenues for further investigation into the cellular functions of Ube2j2 under basal and ER-stressed conditions.

## Introduction

Protein ubiquitylation plays a central role in cell physiology by regulating a vast number of protein activities. The ubiquitin-proteasome system is well-established as a central axis for regulating targeted protein abundance, and it is also crucial in maintaining protein homeostasis (proteostasis) in conjunction with molecular chaperones. The general structure of interactions in the “ubiquitin cascade” utilizes increasing specificity to ensure that substrates are ubiquitylated properly. In mammalian cells, two E1 ubiquitin-activating enzymes are responsible for the ATP-dependent charging of ubiquitin, which is then transferred to one of approximately 40 E2 ubiquitin-conjugating enzymes (E2s) (van Wijk and Timmers 2010; Parashar et al. 2025). Ubiquitin-charged E2s interact directly with unique combinations of the hundreds of E3 ubiquitin-protein ligases (E3s) to transfer ubiquitin onto substrate molecules (Ye and Rape 2009; Stewart et al. 2016; Gundogdu and Walden 2019). Ubiquitylation can also be reversed by deubiquitylating enzymes (DUBs) (Fischer 2003; Sowa et al. 2009). Depending on the number and pattern of ubiquitin modifications, substrate proteins may be re-localized, activated, repressed, or degraded. Thus, specific cohorts of E2 conjugases and E3 ligases can have wide-ranging effects on cellular pathways.

E2 enzymes provide an initial element of subcellular and temporal regulation to the ubiquitylation cascade through spatial restriction (van Wijk et al. 2009; Ye and Rape 2009; van Wijk and Timmers 2010; Witus et al. 2021). E2s with specific subcellular localizations like the endoplasmic reticulum (ER), cytosol, nucleus, or mitochondria, are positioned to interact with E3 ligases to facilitate ubiquitylation of more narrowly defined classes or groups of proteins (Stewart et al. 2016; Christianson and Carvalho 2022). Nonetheless, some E3-containing complexes, such as those that function in ER-associated degradation (ERAD), target diverse groups of substrates (Carvalho et al. 2006; Ravid et al. 2006; Sato et al. 2009; Sharninghausen et al. 2024) despite being spatially constrained.

The ERAD machinery is transcriptionally controlled downstream of the Unfolded Protein Response (UPR), which coordinates transcriptional and translational responses to proteotoxic stress caused by e.g., protein misfolding and aggregation, hypoxia, and oxidative stress (reviewed in Hetz et al., 2015). The overall aim of the UPR is to reinstate cellular homeostasis and, should this fail, initiate apoptosis. ERAD complexes identify and remove misfolded proteins from the ER lumen and membrane and target them for degradation by the proteasome (reviewed in Christianson et al., 2023; Christianson and Carvalho, 2022). This process is dependent on an expanding suite of recognized E3 ligases that partner with ER-localized E2 conjugases.

The ER-localized E2 enzymes Ubc6 and Ubc7 were originally identified in yeast as critical members of the ERAD machinery, and the mammalian orthologs, Ube2j1/Ube2j2 and Ube2g1/Ube2g2 were subsequently identified (Lenk et al. 2002; Wang et al. 2009; Abdul Rehman et al. 2024; Swarnkar et al. 2024). Ubc6 and Ubc7 facilitate E3-dependent ubiquitylation by interacting with the E3s Doa10 and Hrd1, respectively (Kreft and Hochstrasser 2011; Zattas and Hochstrasser 2015; Lips et al. 2020; Mehrtash and Hochstrasser 2022; Christianson et al. 2023). In addition, they participate in sequential ubiquitylation, whereby Ubc6 first catalyzes the addition of ubiquitin to an [atypical] hydroxylated amino acid (Ser or Thr) and Ubc7 then extends a ubiquitin chain via [typical] lysine modification (Weber et al. 2016). Ube2j2, in particular, displays a preference for hydroxylated amino acids (Wang et al. 2009; Abdul Rehman et al. 2024; Swarnkar et al. 2024), whereas Ube2j1 can utilize hydroxylated residues *in vivo* (Burr et al. 2013). Also similar to yeast, Ube2j2 and Ube2g2 interact with distinct ER-localized E3 ligases, including MARCH6 and HRD1, the mammalian orthologs to Doa10 and Hrd1, respectively, along with RNF139 and RNF145 (reviewed in Christianson and Carvalho 2022; Vrentzou et al. 2025). The conserved partnership between ERAD E2s and E3s is an indicator of their central role in cellular function.

Human Ube2j1 and Ube2j2 are similar (43% amino acid sequence similarity, 27% identity), and their similar (though not identical) biochemical activity suggests overlapping functions in the cell. However, investigations into the activities and targets of Ube2j2 in mammalian cells highlight its unique targets and physiological roles (Younger et al. 2006; van den Boomen et al. 2014; van de Weijer et al. 2014; Glaeser et al. 2018; Liu et al. 2019; Cremer et al. 2021; Wolf et al. 2021; Yu et al. 2025). Recently, Ube2j2 has also been shown to contribute to progression and drug resistance in cancer models (Kumari and Kumar 2023; Nakao et al. 2023; Lin et al. 2024). Thus, Ube2j2 plays a distinct role in mammalian protein homeostasis and cell physiology.

To uncover cellular pathways associated with Ube2j2, we first used mass spectrometry to quantify the proteomes of Ube2j2 CRISPR-derived knockout cells under control and tunicamycin-induced ER stress conditions. We then analyzed the proteomic data using a computational network-based procedure that combined the *weighted gene co-expression analysis* (WGCNA) (Langfelder and Horvath, 2008) and differentially expressed proteins (DEP) (Robinson et al. 2010) approaches to model changes in protein abundance and hereby provide a systems-level view of molecular differences between treatments (Serin et al. 2016; Vella et al. 2017). Thus, we were able to (1) identify sets of proteins (modules) with similar abundance profiles across experimental conditions, (2) define module eigengenes (representative profiles that summarize the behavior of any given module, calculated as the first principal component of the protein abundance data) and relate these to experimental treatments, (3) annotate the modules using Gene Ontology (GO) enrichment analysis to extract biological meaning (Ashburner et al. 2000), and (4) find key drivers in the different modules (so-called hub genes) by identifying highly connected proteins that had strong significance in the DEP analysis. To avoid confusion with the term “hub protein” (a central protein in a protein-protein interaction network), we here employ the term “hub gene”, which originates from the WGCNA approach, even though the present analysis is applied to protein abundance. Such hub genes provide potential targets for future study into the biological processes associated with Ub2j2 expression and/or ER stress.

## Results

We first used CRISPR-Cas to create a cell line of U2OS cells, C6β (denoted KO or C6 in raw data sets), lacking a region of 432 base pairs spanning the 5’ UTR and first coding exon of the *UBE2J2* gene and not displaying any visible Ube2j2 expression as probed by Immunoblotting (**Figure S1**). Then, to gain insight into changes in protein abundance as a result of ER-stress duration and the presence or absence of Ube2j2, we grew (in quadruplicate) wild-type and Ube2j2 KO cells in either dimethyl sulfoxide (DMSO; vehicle control) or 10 µg/mL tunicamycin – a cell permeable agent that prevents N-glycosylation in the ER and thereby leads to protein misfolding and activates the UPR – for 6 or 18 hours. Upon flash-freezing, the cells were then processed for and analyzed by quantitative mass spectrometry.

The resulting 32 data sets contained 5903 unique protein sequences, and the quality of samples was gauged based on the coefficient of variation of replicates, which ranged from 11.24 to 18.77% within a sample group. Samples with a coefficient of variation of 10-20% are generally considered reproducible for downstream analyses (Cho et al. 2021). The WGCNA approach defined 12 statistically separate protein abundance profiles each comprising 200-1029 proteins (**Figure 1a, Supplementary File 1**). These 12 modules were annotated to describe the expression and functional properties of the proteins within each module and their association to experimental factors. As visualized by the eigengene values for each module (**Figure 1b**), variation in protein levels across genotypes was the primary factor in defining modules, followed by tunicamycin treatment, and to a lesser extent time (**Figure 1c, Supplementary File 2**). Eigengenes represent the absolute value of changes in abundance, so the magnitude of the change, rather than the direction of change, in protein abundance is reflected in the eigengene pattern. The identified modules also had significant enrichment for specific GO categories in biological processes, cellular components and to a lesser extent molecular functions (**Supplementary File 3**), allowing additional insight into the functional properties of modules.

**Figure 1.**
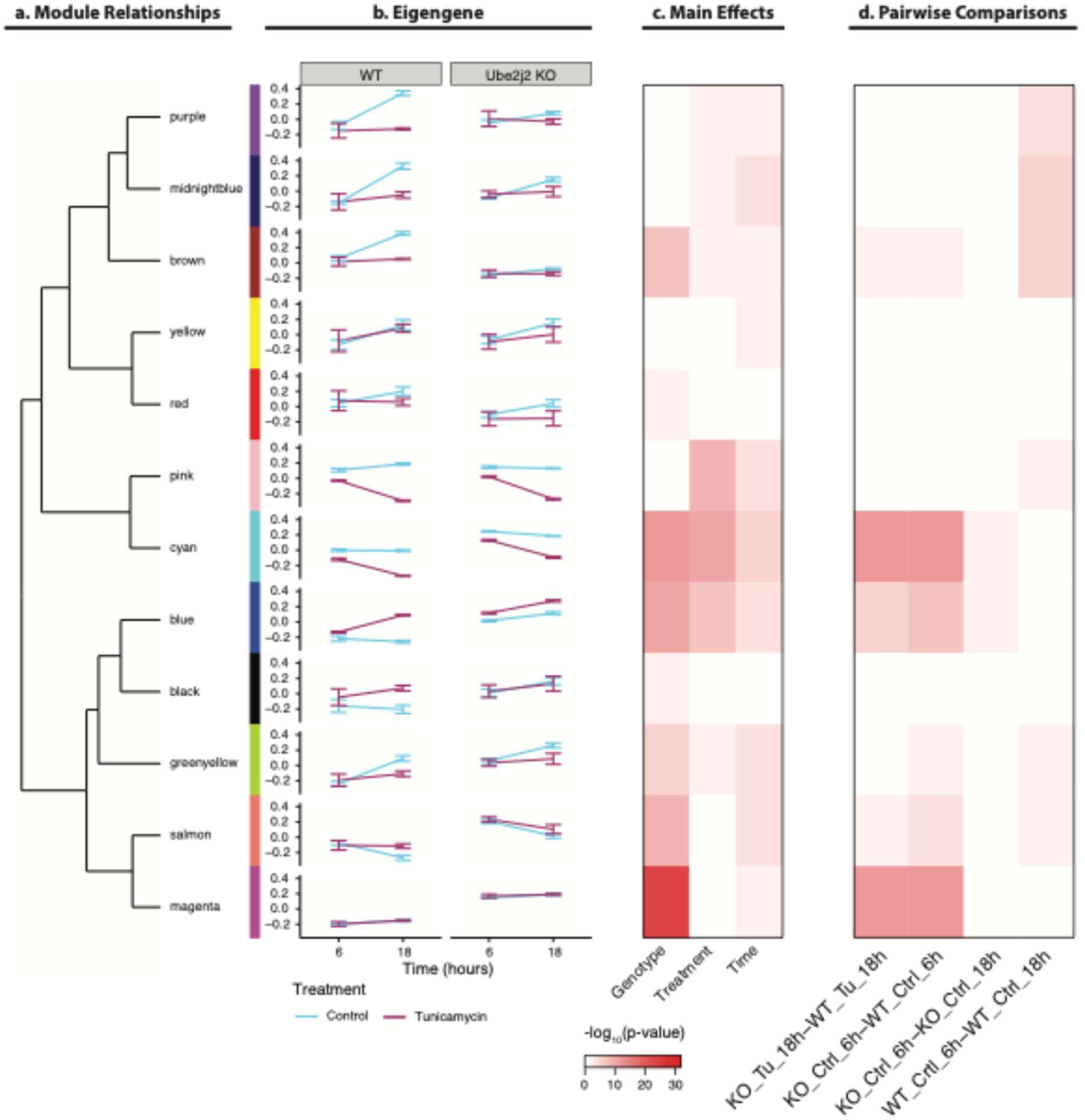
Overview of proteomic data gathered from WT and Ube2j2 knockout (KO) cells in four conditions. (**a**) Dendrogram showing the relatedness of eigengenes between modules (named by color). (**b**) Line graphs of eigengene expression from the wildtype (WT) and Ube2j2 mutant (KO) genotypes across treatment (control and tunicamycin) and time (6 and 18 hours). (**c**) Heatmap of –log10 p-value of module eigengene significance for main effects from an ANOVA model. (**d**) Heatmap of –log10 p-value of the comparisons of interest for each of the modules. The smaller the p-value (the warmer the color), the more significant the difference between conditions. ALT TEXT: Graphical figure with panels labeled a through d. The figure includes a dendrogram that shows the relationship between different protein abundance modules (named by colors). The eigen gene that describes the changes in protein abundance in wildtype and Ube2j2 cells lines is shown next to each module. Two heat maps show the statistical relevance of different conditions and pairwise comparisons between conditions for each module with darker colors representing lower p-values.

Four pairwise contrasts were selected to understand the effects that tunicamycin treatment has on the Ube2j2 KO cells: KO_Tu_18hr-vs-WT_Tu_18hr; KO_Crtl_18hr-vs-WT_Ctrl_18hr; KO_ Crtl _6hr-vs-KO_Ctrl_18hr; WT_ Crtl _6hr-vs-WT_ Crtl _18hr. These four comparisons were chosen for their potential to provide insight into differences in protein abundance based on changes of a single variable at a time (genotype, drug treatment, or time in culture). We chose to focus on treatment conditions at 18 hours, since this late timepoint was expected to provide maximal changes in protein abundance. Further dissection of the modules showed that 9 out of the 12 modules were associated with at least one of these selected pairwise contrasts of interest (**Figure 1d, Supplementary File 2**). Below, we focus on four modules or groups of modules whose patterns of protein abundance levels support our current understanding of the functions of Ube2j2 in protein homeostasis at the ER and offer new insights into cellular protein networks associated with the loss of Ube2j2 and/or ER stress.

### Pink module: Response to tunicamycin is not affected by loss of Ube2j2

The pink module includes proteins whose abundance is affected strongly by tunicamycin-induced stress but not by cellular genotype. The eigengene for the pink module shows stable abundance over time and between the wild-type and knockout cells (**Figure 1b**). However, treatment with tunicamycin leads to a strong up or down regulation of these proteins, relative to the control, regardless of genetic background. Proteins included in the pink module represent well-characterized members of ER-stress pathways such as HERP, HSPA5 (BiP), and VIMP (SelS) (**Figure 2a-c)**, all of which also showed clear upregulation in response to tunicamycin via immunoblotting (**Figure 2d)**. In this manner, the pink module provides an important validation of our mass spectrometry data. **Figure 2e** shows the distribution of proteins in the pink module annotated by biological process ontology. These are enriched for protein processing, folding, and homeostasis as well as protein synthesis and metabolism. Processes related to cytokinesis and chromosome segregation are also identified in this module.

**Figure 2.**
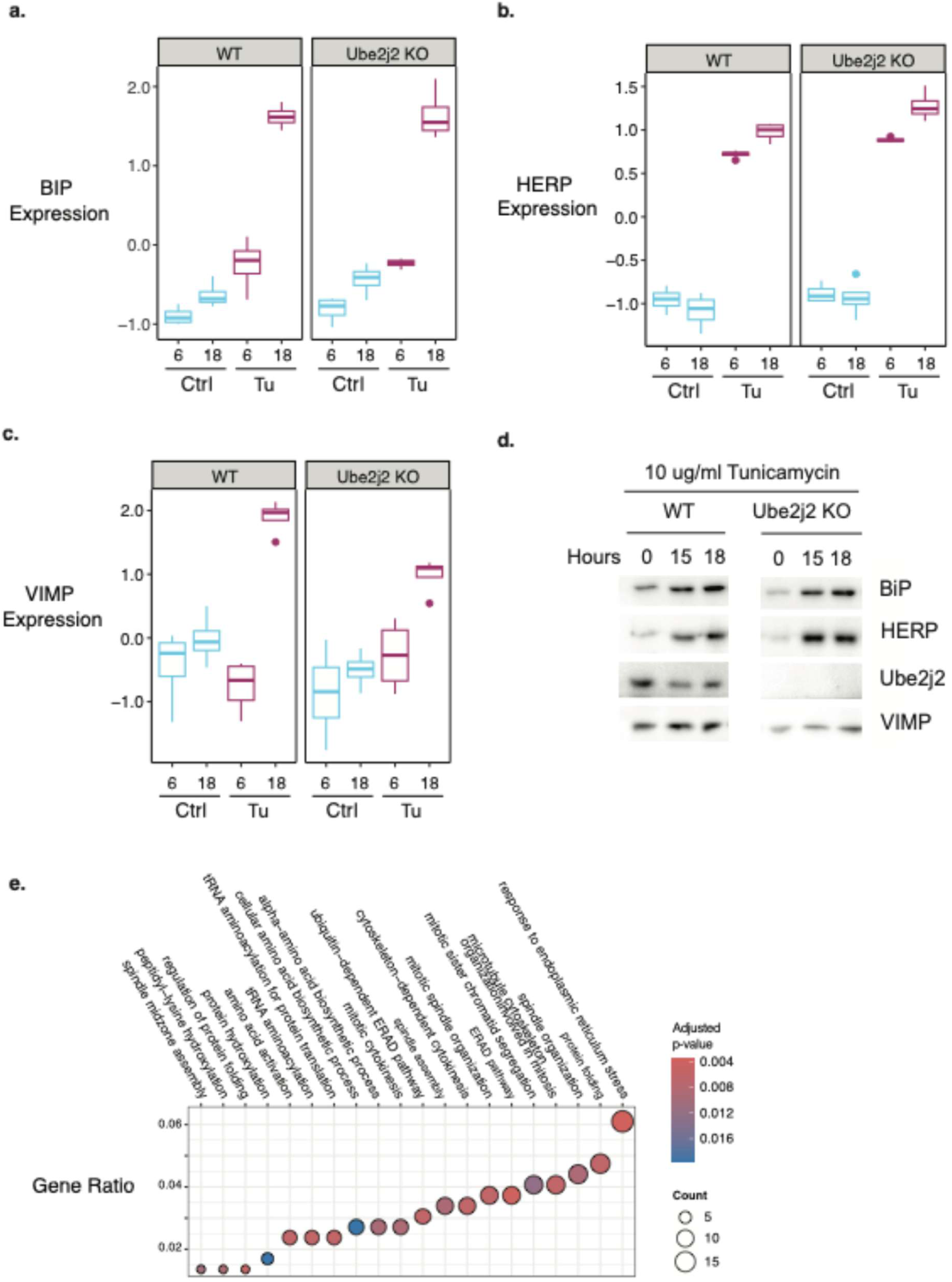
Quantitative relative expression data and immunoblot validation for individual proteins in the pink module. Graphs show changes in protein abundance for (**a**) BiP, (**b**) HERP, and (**c**) VIMP in wildtype and Ube2j2 deletion cells in each of the four conditions (DMSO control at 6 and 18 hours, or tunicamycin treatment at 6 and 18 hours). (**d**) Representative immunoblot of U2OS cells following treatment with tunicamycin for the indicated times. Cell lysates were normalized by a BCA assay prior to SDS-PAGE loading, and blots were probed for the three UPR-associated proteins and Ube2j2. (**e**) GO enrichment analysis of proteins present in the pink module using the biological processes ontology. The graph shows the top 20 most significant terms (<u>adjusted p-value of <0.05)</u> and the count of module proteins observed in each term. ALT TEXT: Three box-and-whisker plots labeled a through c show how the abundance of three individual proteins change in wildtype and Ube2j2 knockout cells under different conditions. An immunoblot labeled d shows corresponding biochemical data for the same three proteins.

### Magenta module: Loss of Ube2j2, regardless of treatment with tunicamycin, drives changes in protein abundance

The magenta module includes 1029 proteins whose abundance is dependent on the presence or absence of Ube2j2 (p-value < 0.001), with little-to-no additional effect of tunicamycin or time (**Figure 1b**). Functional analysis of the magenta module identified significant associations with terms from the cellular components and biological processes, and to a lesser extent molecular function ontologies (**Supplementary File 2).** Based on this analysis, the magenta module was enriched with three main categories (**Figures 3a and 3b**). First, actin-binding proteins, including some that are involved with vesicular transport and exocytosis; second, regulation of mitotic cell cycle involving spindle localization and segregation of sister chromatids; and third, processes that are important for protein abundance regulation and homeostasis. To further narrow down the list of proteins in the magenta module, we conducted a hub gene analysis and identified 135 proteins that were highly connected in the magenta module and significantly associated with protein abundance differences between Ube2j2 KO versus wildtype cell lines (**Figure 3c**). GO enrichment analysis of magenta hub genes by cellular component (**Figure 3d**) found similar results to the module level analysis and suggest that hub genes play a critical role in defining the functions associated with actin and the regulation of the proteasome. Interestingly, Ube2g2, the closely related E2 enzyme that works in tandem with Ube2j2 in yeast (Weber et al. 2016; Mehrtash and Hochstrasser 2022) is also included in this module. This suggests that at least some of the proteins in the magenta module may directly compensate for the loss of Ube2j2’s enzymatic activity. Thus, at least two major cellular components – actin-based motility and cellular structure, and protein degradation – are influenced by the presence or absence of Ube2j2, regardless of tunicamycin-induced ER stress.

**Figure 3.**
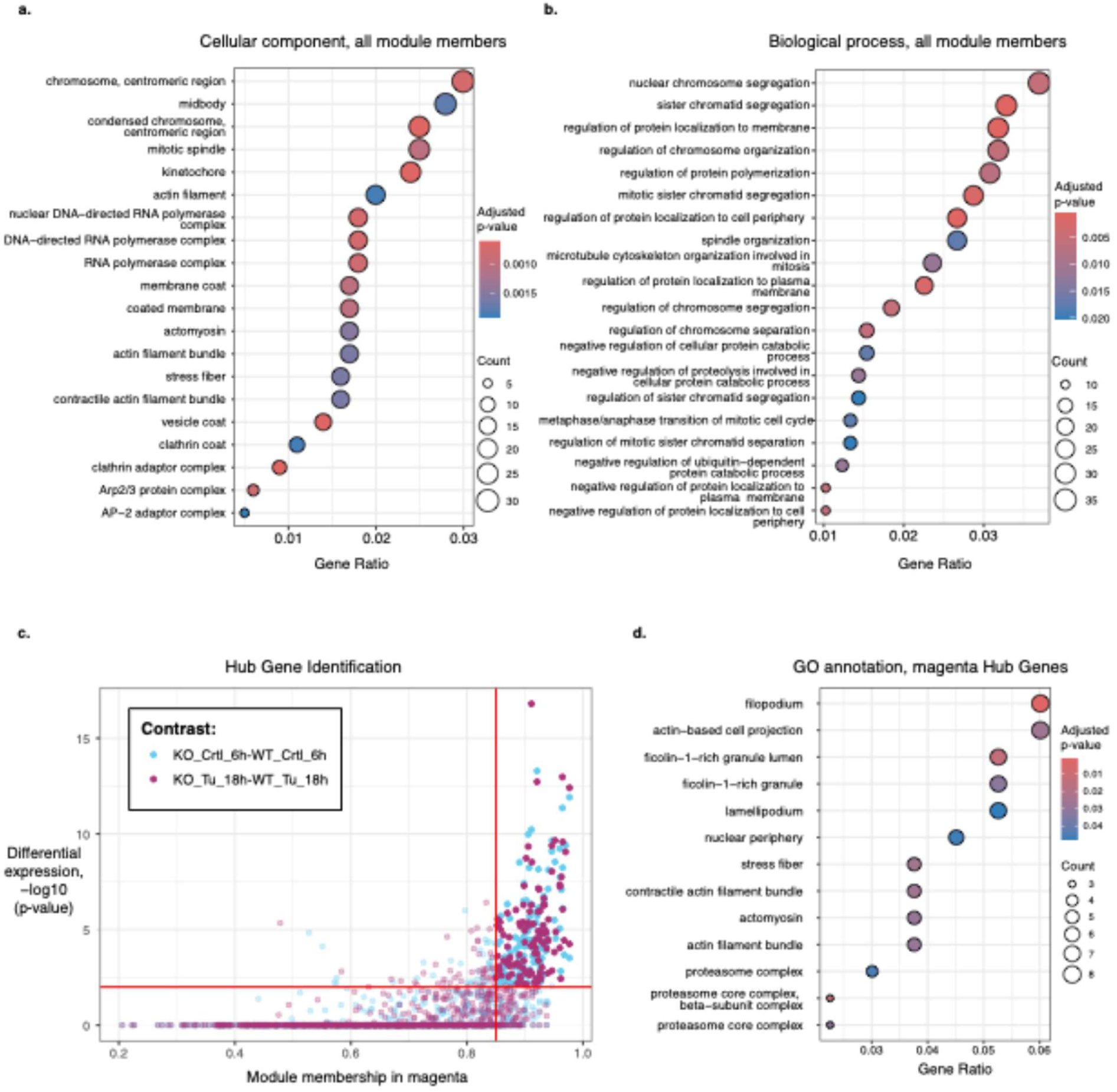
Functional annotation and hub gene analysis for the magenta module. GO enrichment analysis of all protein in the magenta module using the (**a**) cellular component and (**b**) biological processes ontology. The graphs show the most significant terms (adjusted p-value of <0.05) and the count of module proteins observed in each term. (**c**) Hub genes (upper right quadrant) in the magenta module were identified as genes having both a module membership greater then 0.85 and significant (adjusted p-value <0.05) differential abundance with at least one of the pairwise comparisons that relate to changes in protein abundance between KO vs WT cells over time. Proteins that are in the magenta module but are not hub genes are displayed in muted colors in the remaining three quadrants of the graph. (**d**) GO enrichment analysis of all proteins present in the magenta module using the cellular component ontology. The graph shows the top 20 most significant terms (adjusted p-value of <0.05) and the count of module proteins observed in each term. ALT TEXT: Three graphs (a, b, d) show the distribution of GO annotations across proteins and hub genes in the magenta module. Circles representing the number of proteins with any given GO annotation are colored according to their statistical significance, with red and blue representing higher and low statistical significance, respectively. Graph c is divided into quadrants. The upper right quadrant includes subset of hub genes in the magenta module accumulate differently depending on their exposure to tunicamycin in Ube2j2 cells.

### Blue and black modules: Loss of Ube2j2 phenocopies treatment with tunicamycin

The blue and black modules include proteins whose abundance in response to tunicamycin treatment is phenocopied by the loss of Ube2j2 (**Figure 1b)**. Proteins that are members of the black and blue modules are affiliated with a broad set of biological processes including protein-complex interactions, translation initiation, and protein and RNA-localization (black module, **Figure 4a**) and vesicular and ER-to-Golgi transport, electron transport, and nucleotide metabolism (blue module, **Figure 4b**).

**Figure 4.**
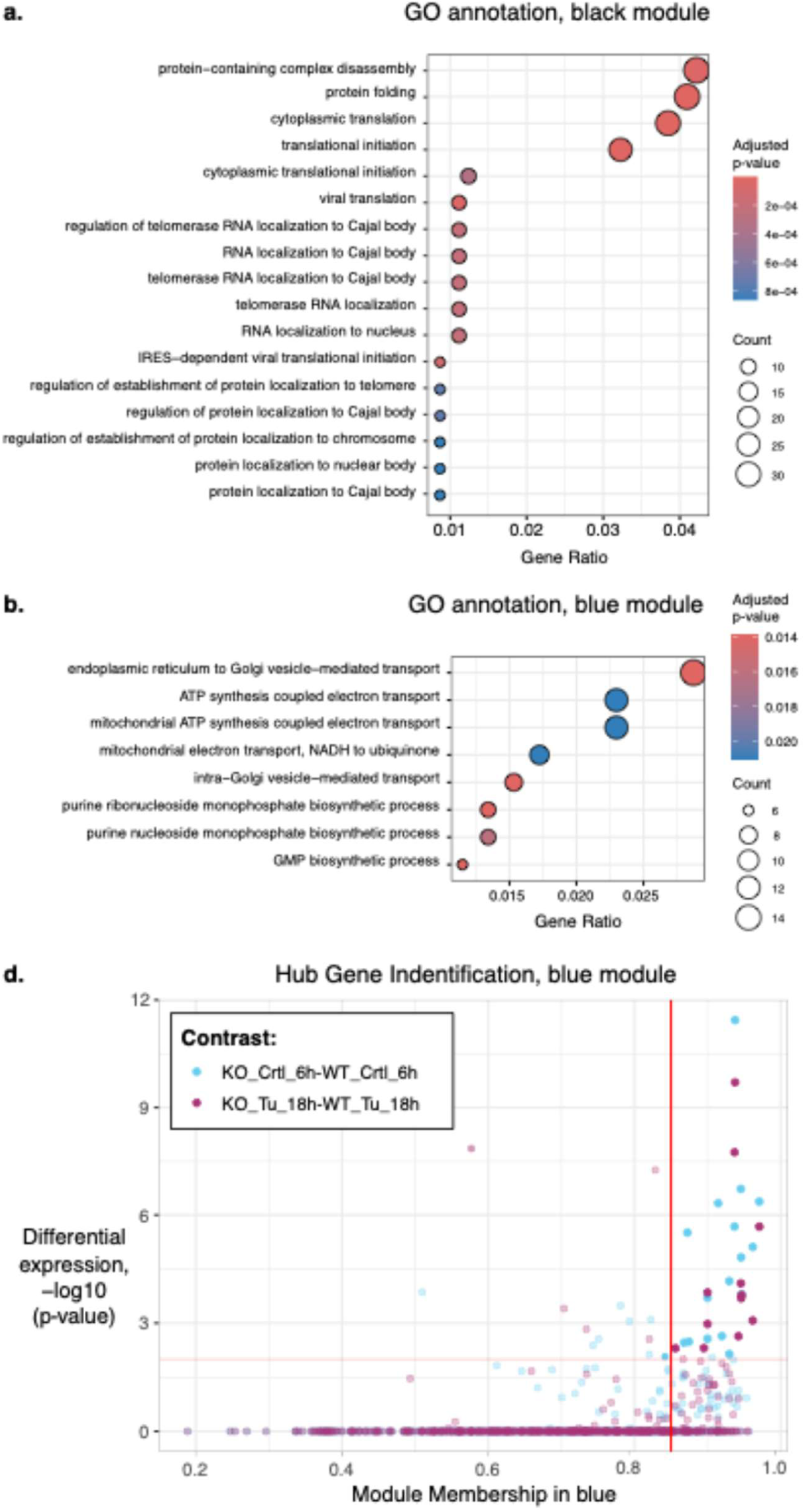
Analysis of the black and blue modules. (**a)** GO enrichment of proteins present in the black module using the biological process **(b**) GO enrichment of proteins present in the blue module using the biological process. Graphs show the top significant (adjusted p-value < 0.05) terms and the count of module proteins observed in each term. (**c**) Identification of hub genes (upper right quadrant) in the blue module showing proteins with high module membership (>0.85) and significant association with at least one pairwise comparison that relate to changes in protein abundance between KO vs WT cells over time. ALT TEXT: Two graphs (a, b) show the distribution of GO annotations across proteins and hub genes in the black and blue modules, respectively. Circles representing the number of proteins with any given GO annotation are colored according to their statistical significance, with red and blue representing higher and low statistical significance, respectively. Graph c is divided into quadrants. The upper right quadrant includes subset of hub genes in the blue module accumulate differently depending on their exposure to tunicamycin in Ube2j2 cells.

While there are no identified hub genes for the black module, the blue module includes 20 hub genes (**Figure 4c)**. Among these, a cluster of COPI coatomer proteins – COPB1, COPB2, COPG1, and COPG2 – are striking due to their roles in Golgi-to-ER transport. In addition, protein homeostasis components including the proteasome adapter and scaffold protein ECM29, UFM1- and SUMO-activating proteins (UBA5 and SAE2, respectively), and the E3 ubiquitin ligase, listerin, are blue module hub genes, along with the translation initiation factor EIF2A, which plays a central role in regulating the PERK branch of the UPR (Harding et al. 2000; Hetz et al. 2015b). Thus, many blue module hub genes have roles in proteostasis. Only four of the 20 hub genes for the blue module have significant differences between 18 hours of tunicamycin treatment in wildtype cells and 6 hours of basal conditions in mutant cells. Overall, we find that the blue and black modules describe proteins whose abundance in the absence of Ube2j2 is similar to that of ER stress conditions, and for which additional ER stress does not amplify the effect of the loss of Ube2j2.

### Purple, midnightblue, and brown modules: Effect of time in culture is abrogated in the absence of Ube2j2

The three related modules purple, midnightblue, and brown describe a pattern wherein time-dependent changes in protein abundance under control conditions are attenuated when Ube2j2 is missing (**Figure 1b**). This could represent a change in cell-cycle progression or disparate reactions to time in culture.

In all three modules, proteins involved with ribosomal metabolism (including biogenesis and rRNA processing) are highly represented (**Figure 5**), suggesting a previously unappreciated role for Ube2j2 in ribosome biosynthesis and regulation. The GO annotations of proteins in the purple, midnightblue, and brown modules suggest that Ube2j2 may also impact protein translation during cellular growth in culture, or potentially during advancement of the cell cycle (**Figure 5**).

**Figure 5.**
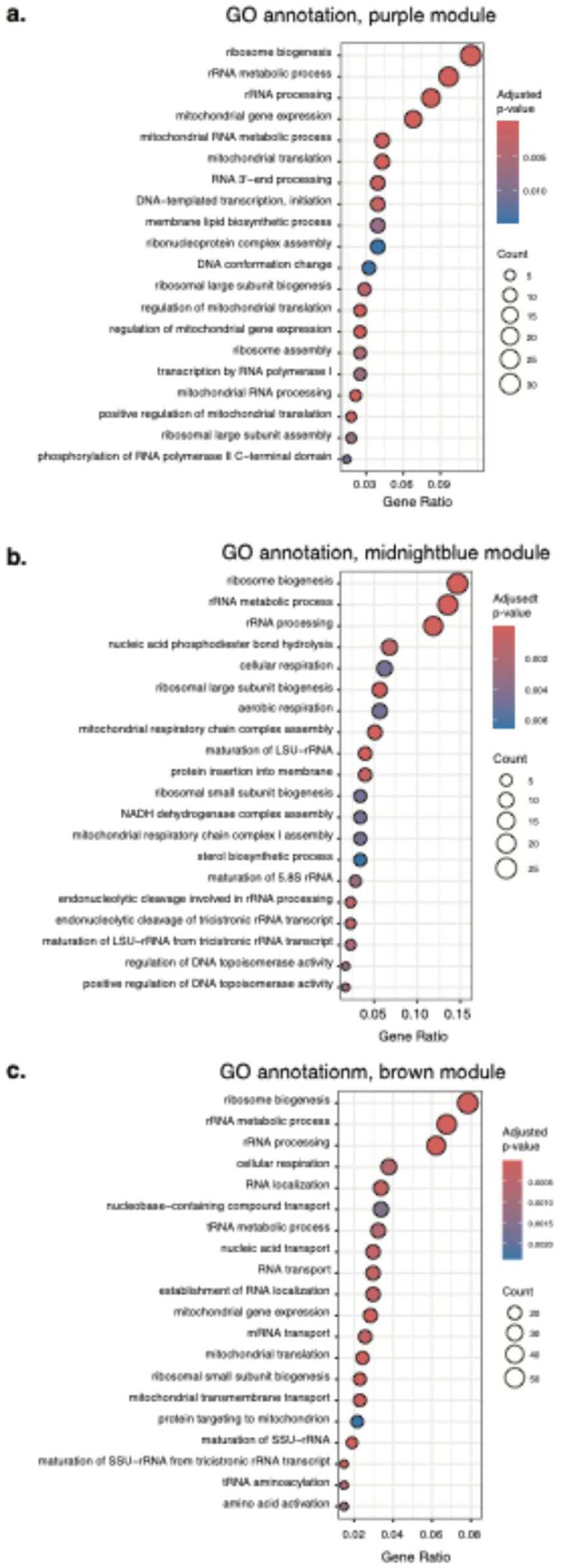
GO enrichment analysis of proteins present in the (**a**) purple, (**b**) midnightblue, and (**c**) brown modules using the biological process ontology. Terms are significant with an adjusted p-value of <0.05. ALT TEXT: Three graphs labeled a, b, and c show the distribution of GO annotations across proteins and hub genes in the purple, midnightblue, and brown modules. Circles representing the number of proteins with any given GO annotation are colored according to their statistical significance, with red and blue representing higher and low statistical significance, respectively.

Only the brown module has associated hub genes and about one third of these 23 hub genes are directly associated with the ribosomal metabolism described by the annotation of the entire module. This suggests that those proteins will be strong candidates for future research into the role of Ube2j2 in regulating cellular health during culturing and growth.

## Discussion

Here, we engineered a line of U2OS cells lacking Ube2j2 and profiled their proteome alongside wild-type cells under four different conditions. Our analysis resulted in 12 modules that describe patterns of changes in protein abundance across nearly 6000 protein sequences. We identified four modules/groups of modules that are characterized by interactions between cellular genotype, treatment condition, or time in culture. GO annotation of the proteins that most strongly drive the eigengene pattern for these modules (hub genes) show that defined classes of biological functions are associated with the magenta, black and blue modules, and that three closely related modules (brown, midnightblue, and purple) show overlapping representation of the same or similar biological function.

There are, to our knowledge, two other proteomic datasets available for cells lacking wildtype Ube2j2 levels. One investigation showed that differentially downregulated proteins in Ube2j2 knockout testes cells were enriched in gamete-specific proteins (Yu et al. 2025). The other investigation, a large-scale transcriptomic and proteomic study, found that the combined knockdown of Ube2j2 and the E1 ligase UBA1 reduced K48-linked ubiquitylation and resulted in the concomitant downregulation of Ube2D1 and Ube2E3 mRNA levels and Ube2D1 and Ube2E1 protein levels in HEK293T cells (Hunt et al. 2023). The present data set is unique because it probes the response of cells to the stress that Ube2j2 would normally help to alleviate.

Our dataset and analysis add to previously published data sets by combining the complete knockout background with a network-based perspective on changes to the proteome in response to two variables, time and drug-induced stress. Our analyses provide a suite of protein-abundance profiles that reveal both expected and unpredicted changes across the proteome in the absence of Ube2j2. Our results highlight UPR and ubiquitin-dependent protein homeostasis networks (highly represented in the pink module and magenta modules (Figures 2 and 3)). The network analysis also suggests Golgi-to-ER trafficking pathways (magenta, blue, and black modules, Figures 3 and 4) and RNA metabolism pathways (brown, purple, and midnightblue modules, Figure 5) that are dependent on Ube2j2. In addition, we were interested in any changes in abundance of the Ube2j2 paralog, Ube2j1, and the two Ubc7 homologs, Ube2g1 and Ube2g2 in this dataset. Ube2j1 and Ube2g1 are found in the brown module, which makes them members of a cohort of proteins whose reaction to time in culture is modulated by Ube2j2 (Figure 1), while Ube2g2 is a hub gene for the magenta module (Figure 3). These results suggest different types of compensation by ER-associated E2 enzymes in the absence of Ube2j2. Together, the WGCNA analysis, eigengene profiles, and GO annotations that accompany our data sets suggest inroads for future research by providing insights into changes in protein abundance that are based entirely on the presence or absence of Ube2j2 or through complex interactions between Ube2j2 and tunicamycin-induced stress and/or time in culture.

## Materials and methods

### Generation of CRISPR-Cas9–Mediated UBE2J2 knockout cell line

Human osteosarcoma (U2OS) cells were subjected to two rounds of CRISPR-mediated gene editing. Cells were transiently co-transfected with plasmids encoding Cas9 and single guide (sg) RNAs targeting UBE2J2 (sgRNA#3: 5′-CAGTAAGAGGGCTCCGACCA-3′ and sgRNA#1: 5′-CCGGGTCTTTCTTAATGCGA-3′) using polyethylenimine transfection reagent. sgRNA sequences were designed and kindly provided by Dr. Dick van den Boomen in the lab Prof. Paul J. Lehner, University of Cambridge, UK. After 24 hours, cells were selected with 10 µg/µl Puromycin and 100 µg/µl Hygromycin B to enrich for transfected cells. To enhance selection efficiency and avoid false positives, cells were split three days post-transfection. Selection medium was replaced once non-transfected controls were eliminated. Due to variable sgRNA efficiency and to obtain a homogenous population of cells, single-cell clones were isolated by limiting dilution (0.3 cells/well in 96-well plates), expanded, and harvested.

Genomic DNA was extracted from potential knockout clones using QuickExtract™ DNA Extraction Solution 1.0 (Epicentre), following the manufacturer’s instructions. Briefly, ∼9,000 cells were pelleted at 500 × g for 5 min, washed in phosphate buffered saline (PBS), and resuspended in 50 µL QuickExtract™. Extracts were incubated at 70°C for 30 minutes with intermittent vortexing, followed by 95°C for 5 min.

The extracted genomic DNA was used as template for PCR amplification prior to TOPO cloning and Sanger sequencing. PCR reactions were performed using the Phusion DNA polymerase (Thermo Fisher Scientific) with a primer pair designed to amplify a region of the gene covering the sgRNA target site (FWD: 5′-GGCCAGGGCATAAGAGTTTATTTAC-3′, REV: 5′-TACTGCTCTGTCGAAACACTGC-3′), yielding an expected amplicon size of 1472 bp. PCR products were verified by agarose gel electrophoresis and subsequently ligated into the pCR™-Blunt II-TOPO™ vector using the Zero Blunt™ TOPO™ PCR Cloning Kit (Thermo Fisher Scientific), following the manufacturer’s protocol. The ligation products were transformed into competent *E. coli* DH5α cells, and ten colonies per reaction were picked for overnight culture. Plasmid DNA was isolated using mini-prep procedures and subjected to Sanger sequencing (Eurofins) using the M13 reverse −29 primer (5′-CAGGAAACAGCTATGACC-3′). Sequencing analysis revealed a 432 bp deletion in the targeted *UBE2J2* locus (**Supplementary Figure 1**).

### Generation of a monoclonal anti-VIMP antibody

Mouse monoclonal antibodies were raised to human VIMP using a previously described protocol (Hendil 2005). For immunization, we used the His_6_-lipoFF-tagged cytosolic region (residues 49–189) of human VIMP with the Sec residue at position 188 replaced with a Cys residue. The protein was produced recombinantly as previously described (Christensen et al. 2012).

### Cell culture, tunicamycin treatment, and harvest

Wild-type or knock-out U2OS cells were grown maintained in Dulbecco’s Modified Eagles Medium supplemented with L-glutamine and heat-treated fetal bovine serum. Cells were passaged regularly, and experiments were performed with cells that had been passaged fewer than 20 times. For drug treatment experiments prior to immunoblotting, cells were treated by adding 10 µg/mL tunicamycin (final concentration) directly to growth media followed by incubation for the indicated times. As a control, 0.01% DMSO (vehicle control, final concentration) was added to cells, which were then incubated for the longest incubation time.

### Protein concentration determination, SDS-PAGE, and immunoblotting

Cells were harvested by placing plates on ice and washing with PBS, followed by lysis on ice in 40 uL radioimmunoprecipitation assay (RIPA) buffer supplemented with phenylmethylsulfonyl fluoride (0.01%) and cOmpleteMini protease inhibitor mixture (1 tablet per 10 mL of buffer, according to manufacturer’s directions). Cells were scraped off from the plates and sonicated for five seconds on ice. Cell lysates were centrifuged for 7.5 minutes at 12,500 rpm at 4°C and supernatant was removed to fresh tubes for analysis. Protein content of cleared lysates was determined by quantitative Bicinchoninic Acid (BCA, Sigma Aldrich) anlysis in 96-well plate format using bovine serum albumin (Sigma Aldrich) as standard. All dilutions for the standard curve and sample measurements were done in triplicates. The BCA was measured at 562 nm using a plate reader (Tecan) and equivalent portions of lysate were run on acrylamide gels and transferred to polyvinylidene fluoride for immunoblotting. Immunostaining was performed with the following primary antibodies, which were diluted in Tris buffered saline plus 0.05% Tween containing 5% non-fat milk or bovine serum albumin: rabbit anti-BiP/GRP78 (Sigma, GL-19; RRID:AB_477030) (1:1000), rabbit anti-HERP (kind gift from L. Hendershot) (1:1000), rabbit anti-Ube2j2 (kind gift from L. Hendershot) (1:1000), mouse anti-VIMP (described above) (1:1000). Primary antibodies were detected with secondary antibodies conjugated to Horseradish Peroxidase (Pierce) diluted 1:100,000 in Tris buffered saline plus 0.05% Tween containing 5% non-fat milk. Blots were visualized using Pierce Femto ECL according to manufacturer’s directions.

### Cell treatment, growth and harvest for mass spectrometry

Wild-type or knockout cells were grown to approximately 50% confluency in 10 cm plates. Cells were grown as described above and then treated by replacing original growth media with media supplemented with 0.01% DMSO (vehicle control) or 10 µg/mL tunicamycin in DMSO followed by incubation for 6 or 18 hours. Media was removed by aspiration and plates were moved onto ice. Cells were scraped off plates into PBS supplemented with protease inhibitor (cOmpleteMini protease inhibitor mixture was dissolved in PBS according to manufacturer’s direction). Cells were spun gently (5 min at 2000 rpm, at 4°C) and washed twice with ice cold PBS supplemented with protease inhibitors. Cell pellets were snap frozen in liquid nitrogen and stored at −80°C.

### Mass spectrometry procedure

In preparation for mass spectrometry, cell pellets were lysed in lysis buffer (1% sodium deoxycholate, 100 mM Tris-HCl, pH 8.5) and incubated for 10 min at 99°C, followed by sonication in a Bioruptor Pico (20 cycles, 30 s on/30 s off, ultra-low frequency). Lysates were cleared by centrifugation, reduced with 5 mM TCEP for 15 min at 55°C, and alkylated with 20 mM chloroacetamide for 30 min at room temperature. Proteins were digested for 16 h at 37°C using trypsin and LysC at a 1:100 (enzyme:protein) ratio. Peptides were acidified with 1% trifluoroacetic acid and desalted using SDB-RPS StageTips.

Peptides were separated on a 50 cm × 75 µm ID home-packed analytical column containing ReproSil-Pur C18-AQ resin (1.9 µm; Dr. Maisch) using an EASY-nLC 1200 ultra-high-pressure system. Samples were loaded in buffer A (0.1% formic acid) and separated using a non-linear gradient from 5% to 60% buffer B (0.1% formic acid, 80% acetonitrile) over 50 min at a flow rate of 300 nL/min. Peptides were introduced via a CaptiveSpray source with a 10 µm emitter into a timsTOF Pro2 mass spectrometer (Bruker) operated in diaPASEF mode.

MS data were acquired over a 100–1700 m/z range using the standard long-gradient diaPASEF method, comprising 16 diaPASEF scans with two 25 Da windows per ramp, covering a total mass range of 400–1201 Da and an ion mobility range of 1.43–0.6 1/K₀. Collision energy decreased linearly from 59 eV at 1/K₀ = 1.6 to 20 eV at 1/K₀ = 0.6 Vs cm⁻². Both accumulation and PASEF ramp times were set to 100 ms, resulting in a total cycle time of 1.17 s.

Quantification of raw data was performed using Spectronaut v15.5 (Bruderer et al. 2015) in directDIA mode using the *Homo sapiens* UniprotKB FASTA (UP000005640; TaxID 9606). Carbamidomethylation of cysteine was set as a fixed modification; oxidation of methionine and protein N-terminal acetylation were set as variable modifications. Up to two missed cleavages were allowed, and a minimum peptide length of seven amino acids was required. The false discovery rate (FDR) at the PSM, peptide, and protein group level was set to 0.01. The raw data processing was implemented in the automated analysis pipeline of the Clinical Knowledge Graph (Santos et al. 2022) and exported as label-free quantitative (LFQ) intensities. The LFQ intensities were normalized by log2 transformation and proteins with less than at least 2 valid values in at least one group were filtered out. Missing values were imputed using the MinProb approach (width=0.2 and shift=1.8, (Lazar et al. 2016)) and with Mixed imputation.

### Bioinformatics analysis

The quantifications from 5903 proteins were used to identify differentially expressed proteins and generate weighted gene co-expression network analysis.

The differential expression analysis was done using edgeR (v3.40.2; (Chen et al. 2025) and the protein intensities were normalized with the trimmed mean of M-values method. Generalized linear models were used to determine if differences in protein intensities were associated with the main effects from the experimental design: genotypes (wildtype and Ube2j2 KO) and cell treatments (vehicle control and tunicamycin) at 6 or 18 hours. Pairwise contrasts for DEPs were specified using the emmeans package (v1.8.7; Lenth and Piaskowski 2017) in R and focused on the effect of tunicamycin treatment has on Ube2j2 mutants through time (KO_Tu_18hr-vs-WT_Tu_18hr; KO_Ctrl_18hr-vs-WT_Ctrl_18hr ; KO_Ctrl_6hr-vs-KO_Ctrl_18hr; WT_Ctrl_6hr-vs-WT_Ctrl_18hr). To determine if a protein was significantly differential expressed with main experiential factors or pairwise contrasts a Benjamini-Hochberg adjusted p-value was less than 0.05. DEPs with significant pairwise contrasts were also used as a criterion for hub gene analysis.

Co-expression analysis and module identification were conducted using functions from WGCNA (v1.72-1; (Langfelder et al. 2008). The soft threshold was determined as the value producing a >90% model fit to the scale-free topology and low mean connectivity. Co-expression relationships were calculated using unsigned Pearson’s correlation coefficients raised to the soft threshold and grouped using hierarchical clustering of dissimilarity among the unsigned topological overlap matrix. Modules were identified using dynamicTreeCut (v1.62-1; (Langfelder et al. 2008)) with parameters of minimum module size of 100, deep split value of 2, and cut height of 0.99. Module eigengenes were calculated as the first principal component of the expression matrix and modules with correlated (>0.80) eigengene values were collapsed because these modules contained proteins that strongly covary in abundance across samples.

The biological meaning of the co-expression modules was investigated using three approaches. First, we tested if module eigengenes were associated with the main effects of the experimental conditions using generalized linear models. Specifically, we tested if module eigengenes were significantly associated with differences between genotypes (wildtype and Ube2j2 KO) or cell treatments (vehicle control and tunicamycin) at 6 or 18 hours. Second, functional annotation of the modules as conducted using gene ontology (GO) enrichment analysis. GO enrichment for each ontology (biological rocess, molecular function, or cellular component) was performed using the clusterProfiler package (v4.17.0; (Yu et al. 2012) and bioconductor Human annotation database (org.Hs.eg.db; v3.18). Visualization of GO enrichment results were done using enrichplot (v1.29.1) and simplifyEnrichment (v1.6.1; (Gu and Hübschmann, 2023). GO terms were determined to be significant if the Benjamini-Hochberg adjusted p-value was less than 0.05. Third, hub genes within modules were identified by examining the interaction between module membership and the –log10 p-values for at least one of the specific pairwise contrasts. Module membership was calculated as the absolute of the Pearson’s correlation between the module eigengene and protein LFQ intensities. The threshold for module membership was having a greater than 0.85 correlation, and the DEP threshold was a –log10 adjusted p-value of greater than 2.

## Data availability

Cell lines are available upon request. Please refer to the Reagent Table for futher details about cell lines and antibodies. Supplemental files available at FigShare. Supplementary File 1 contains the module assignments and accession numbers for proteins in the dataset. Supplementary File 2 contains the raw and adjusted pvalues used to generate Figure 1C & 1D in this manuscript. Supplementary File 3 contains the GO annotations for proteins in each module. Code generated for the analyses described can be found at: https://github.com/mzinkgraf/Ube2j2. The mass spectrometry proteomics data have been deposited to the ProteomeXchange Consortium via the PRIDE partner repository (Perez-Riverol et al. 2025) with the dataset identifier PXD076153.

## Acknowledgements

We acknowledge Dick J. H. van den Boomen (Cambridge Institute for Medical Research, University of Cambridge, UK) for the design of sgRNA sequences and Linda M. Hendershot (St. Jude Children’s Research Hospital, Memphis, USA) for kindly providing us with rabbit anti-HERP and rabbit anti-Ube2j2 antibodies.

## Funding

Mass spectrometry-based proteomics analyses were performed by the Proteomics Research Infrastructure (PRI) at the University of Copenhagen (UCPH), supported by the Novo Nordisk Foundation (NNF) (grant agreement number NNF19SA0059305). CLD was supported by funding from the American Scandinavian Foundation and the Fulbright U.S. Scholar Program. Support for this research was also provided by the Lizzi og Mogens Staal Fond (grant number 2024-1314). LE was supported by the Department of Biology, University of Copenhagen, and a sabbatical grant from the Lundbeck Foundation (grant number R500-2024-1904).

## Conflict of interest

The authors declare no conflicts of interest.

## Author contributions

Conceptualization, C.L.D. and L.E.; Data curation, C.L.D. and M.Z.; Formal Analysis, M.Z.; Funding acquisition, C.L.D. and L.E.; Investigation, C.L.D., S.H.L., C.L.S., Q.G., E.B., R.H.P. ; Project administration, C.L.D. and L.E.; Supervision, L.E.; Visualization, C.L.D., M.Z., and S.H.L.; writing—original draft preparation, C.L.D., M.Z., and L.E.; writing—review and editing, all authors. All authors have read and agreed to the published version of the manuscript.

## Abbreviations

DEP: differentially expressed protein
DMSO: dimethyl sulfoxide
DUB: deubiquitylating enzyme
ER: endoplasmic reticulum
ERAD: ER-associated degradation
GO: gene ontology
KO: knockout
LB: lysogeny broth
LFQ: label-free quantitative
MS: mass spectrometry
PBS: phosphate buffered saline
SDS-PAGE: sodium dodecyl sulfate-polyacrylamide gel electrophoresis
UPR: unfolded protein response
WGCNA: weighted gene co-expression analysis
WT: wildtype

---

**Supplementary Figure 1.**
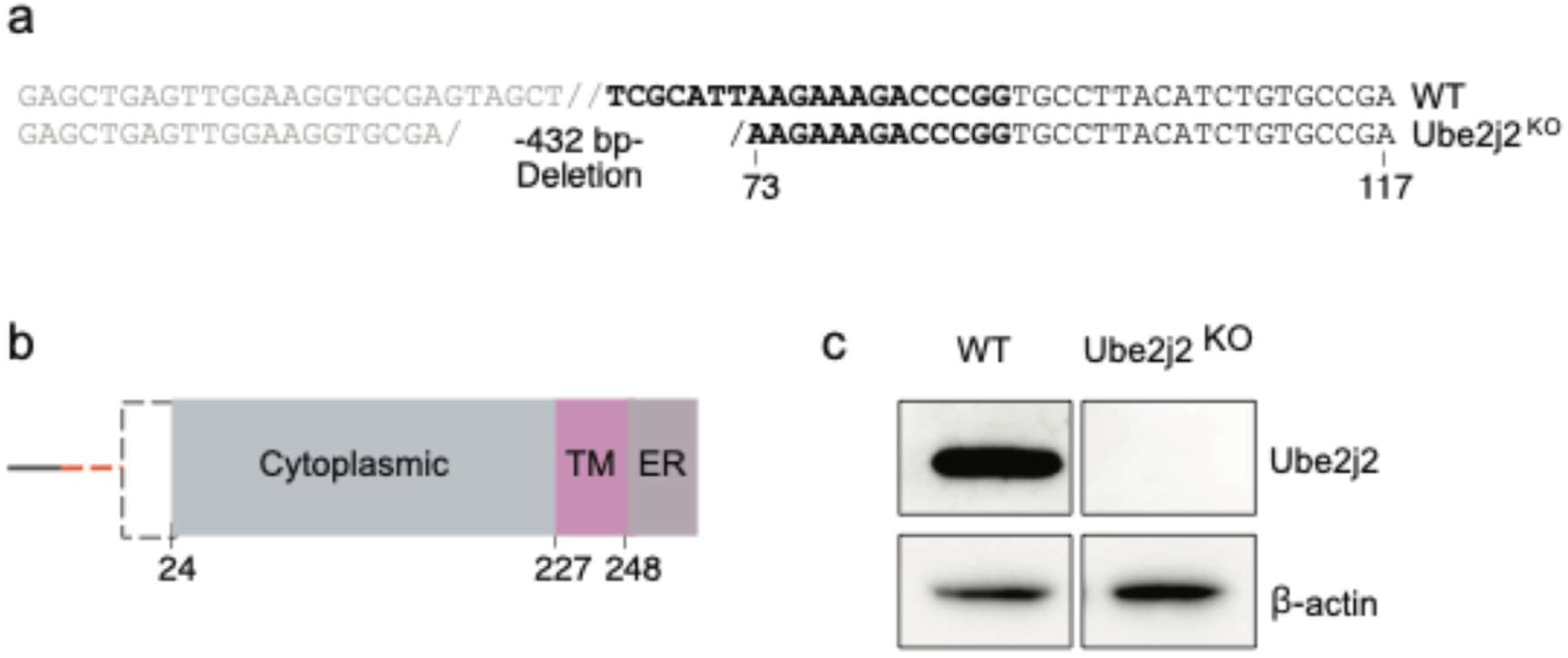
Generation and verification of Ube2j2 KO cell line. (**a**) Nucleotide sequence of the mutated *UBE2J2* clone, illustrating a 432 bp deletion that spans both non-coding (grey) and coding (black) regions. Base numbering is relative to the start of exon 2, the first coding exon. (**b**) Schematic representation of the Ube2j2 protein domain architecture. The deleted region is indicated by black dashed lines in the protein and by an orange dashed line in the upstream nucleic acid sequence. Numbers refer to amino acid residues; TM: transmembrane domain; ER: endoplasmic reticulum lumen localization. (**c**) Immunoblot analysis of protein lysates from wild-type (WT) and *UBE2J2* knockout (KO) U2OS cells. β-actin serves as a loading control. ALT TEXT: A figure with three panels labeled a, b and c. a and c show the DNA sequence and schematic representations of the Ube2j2 knock-out in the cells used in this study. c shows the immunoblot that confirmed the successful removal of the Ube2j2 protein in the cells used in this study.

## Notes

### Competing Interest Statement

The authors have declared no competing interest.

### Summary of Updates

Supplementary data and tables have been uploaded.

